# Distinct cortical populations drive multisensory modulation of segregated auditory sources

**DOI:** 10.1101/2024.12.23.630079

**Authors:** Huaizhen Cai, Harry Shirley, Monty A. Escabí, Yale E. Cohen

**Affiliations:** Department of Otorhinolaryngology: Head and Neck Surgery, University of Pennsylvania School of Medicine, Philadelphia, PA 19104; Departments of Electrical Engineering and Computer Science, University of Pennsylvania School of Medicine, Philadelphia, PA 19104; Biomedical Engineering, University of Connecticut, Storrs, Connecticut; Psychological Sciences, University of Connecticut, Storrs, Connecticut; The Connecticut Institute for Brain and Cognitive Sciences, University of Connecticut, Storrs, Connecticut; Departments of Neuroscience, University of Pennsylvania, Philadelphia, PA; Bioengineering, University of Pennsylvania, Philadelphia, PA

## Abstract

Auditory perception can be modulated by other sensory stimuli. However, we do not fully understand the neural mechanisms that support multisensory behavior. Here, we recorded spiking activity from the primary auditory cortex (A1) in non-human primates, while they detected a target vocalization that was embedded in a background chorus of vocalizations. We found that a congruent video of a monkey eliciting a vocalization improved the monkeys’ behavior, relative to their performance when we only presented a static image of the monkey. As a proxy for the functional organization of A1, we compared the contribution of neurons with significant spectrotemporal response fields (STRFs) with those that had non-significant STRFs (nSTRFs). Based on spike-waveform shape and functional connectivity, STRF and nSTRF neurons appeared to belong to distinct neural populations. Consistent with this, we found that although STRF neurons encoded stimulus information through synchronized activity, the population of nSTRF neurons encoded task-related information in the primate A1 more as a structured dynamic process. Together, these findings demonstrate a functional distinction between the behavioral contributions of nSTRF and STRF neurons.

## Introduction

Our sensory world is inherently multisensory^1,2^. We love wine due to the complex sensory interactions between its taste, aroma, color, and “mouth feel”. Fundamentally, multisensory signals modulate and facilitate perception. The McGurk effect illustrates how visual information (i.e., a speaker’s moving lips) can bias our speech (auditory) perception^3^. Similarly, ventriloquism illustrates that our perception of the spatial location of an auditory stimulus can be biased by visual information^4,5^, whereas the sound-induced flash illusion shows how the number of reported visual events can be biased by auditory information^6^.

Although several studies have examined how spiking activity in the auditory cortex (A1) is modulated by multisensory stimuli^1,2,7–9^, most have not examined A1’s contributions to multisensory behavior, especially in primate models of hearing. Further, of those studies that have examined this relationship, they have been limited in two important ways. First, in general, they have not systematically manipulated multisensory stimuli to test a subject’s psychophysical sensitivity. This is an important manipulation because the relationship between behavioral sensitivity over different psychophysical regimes (e.g., threshold versus suprathreshold) and concomitant changes in neural activity is critical to our understanding of the neural mechanisms of perception^10,11^. Second, previous work often utilized multisensory stimuli in a manner that was not rooted in primate cognition. That is, these stimulus paradigms did not take advantage of the known ways in which the primate brain capitalizes on multisensory signals^2,7,12,13^.

Here, we developed a multisensory task that overcomes these two limitations by training monkeys to detect a target vocalization that was embedded in a background “chorus” of vocalizations. The target was accompanied by either a static video of a monkey with its mouth closed or a dynamic video of a monkey eliciting the target vocalization. We titrated task difficulty by varying the signal-to-noise ratio between the target and the background chorus. While monkeys participated in this ethologically inspired task, we recorded from spiking activity from A1 neurons to identify their contributions to multisensory behavior.

Consistent with previous studies, we found that the dynamic video (relative to the static video) improved the monkeys’ behavior (higher hit rates and decreased decision times) and modified the firing rates of single neurons. We advanced previous findings by demonstrating that A1 neurons with non-significant spectrotemporal response fields (nSTRFs) and those with significant STRFs may originate from distinct neural populations. Further, whereas the population neural trajectories of both STRF and nSTRF neurons encoded the signal-to-noise ratio between the target and background, the trajectories of the nSTRF neurons were modulated more by the visual stimuli than the STRF-neuron trajectories. Thus, nSTRF neurons appeared to play a more direct role in multisensory behavior than STRF neurons. These findings are the first to identify an organizing principle of multisensory information in A1, to characterize a differential functional contribution of STRF and nSTRF A1 neurons to any form of behavior, and to demonstrate that task-related information in the primate A1 is encoded as a structured dynamic process in the neural population.

## Results

We only report data from sessions in which we simultaneously collected behavioral and neural data. Further, we only considered neurons that had stable firing-rate profiles throughout the entire session; in particular, we ensured that the structure of the spectrotemporal response fields and the current-source-density/multiunit-activity profiles were stable before and after the monkeys performed the multisensory task.

### Behavioral performance

Monkeys W (n= 19 sessions) and E (n= 51 sessions) participated in a multisensory task (Fig. 1A) and reliably reported the occurrence of a target vocalization, with performance that depended systematically on the signal-to-noise ratio (SNR) between the target and the background chorus and on the concomitant visual stimulus (Fig. 1B). As the SNR increased, the monkeys’ performance (i.e., hit rate) improved. We saw further performance enhancement when the target vocalization was accompanied by the visually congruent video, relative to the static video (Fig. 1B). On static-visual and congruently visual trials, Monkey E and Monkey W had mean false alarm rates of 5%±1% s.d. and 12%±4% s.d., respectively.

**Figure 1:**
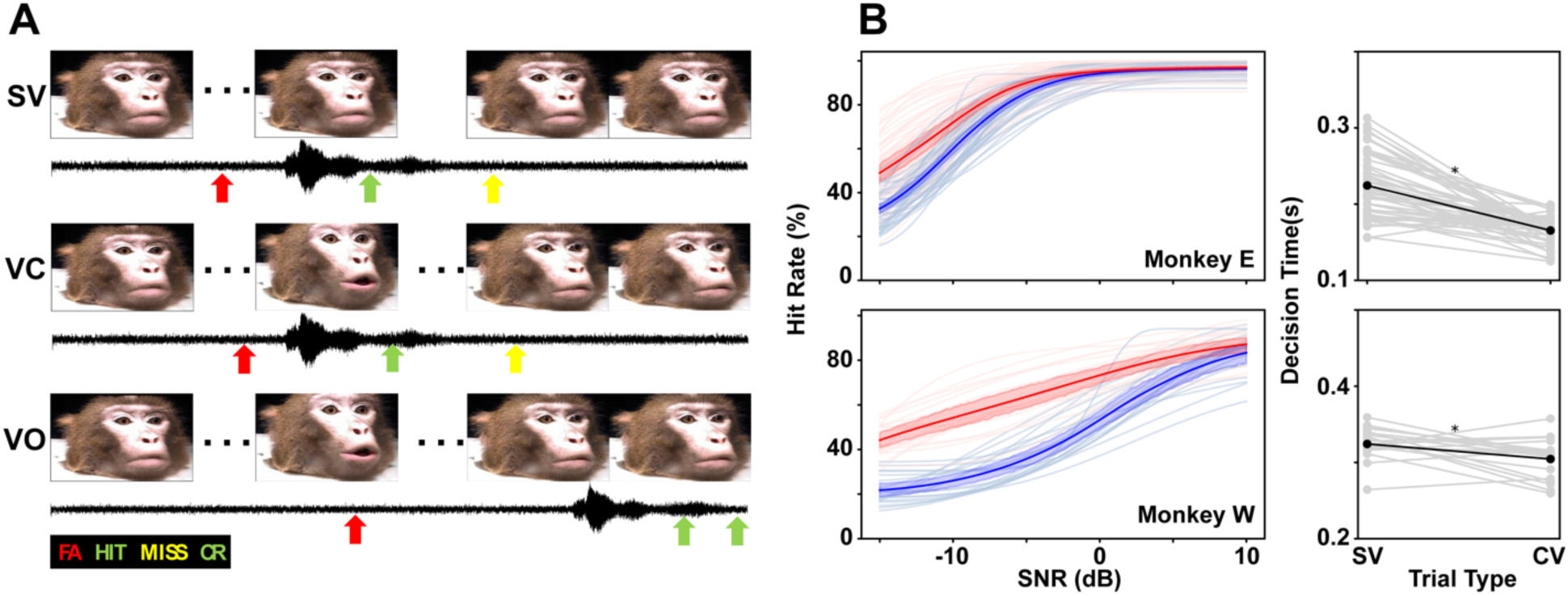
The multisensory task and psychophysical performance. **A.** Monkeys detected a coo vocalization (black acoustic waveform in each panel) in the presence of 1 of 3 different visual-trial conditions: a static-visual trial (SV); a visually congruent trial (VC); and a visual-only trial (VO). In each trial schematic, a red arrow indicates a false alarm, a green arrow indicates a hit trial, and a yellow arrow indicates a miss. In each schematic, a trial unfolds over time from left to right. **B.** Monkeys were more accurate and faster on VC trials than on SV trials. (**left**) Pale lines are psychometric curve fits from individual sessions. Thick lines are fits from the average of the sessions, and the 95% confidence interval is indicated by the shading. The red data are from VC trials, whereas the blue data indicate SV trials. **(right)** Thin grey lines indicate session-by-session decision times (connected by a line across the two visual-trial conditions). The thick black dots are the median decision time.

Using a one-choice drift-diffusion model^14,15^, we calculated the monkeys’ “decision time” during the multisensory task. Unlike the better-known drift-diffusion models that quantify decision times during a two-alternative forced-choice task^16,17^, we instantiated a version that incorporated a single choice (i.e., the detection of the target vocalization). The decision time was the time between onset of the target vocalization and the time to cross the bound (i.e., the decision criterion). We found that decision times were faster (Fig. 1B; Wilcoxon signed-rank test; H_0_: median decision time was the same; Monkey W: p<0.004; Monkey E: p<0.00001) on trials during visually congruent trials (Monkey W: median [interquartile range or IQR] = 0.30 s [0.28–0.32]; Monkey E = 0.17 s [0.15–0.17]) than during static-visual trials (Monkey W: median [IQR] = 0.32 s [0.32–0.33]; Monkey E: median [IQR] = 0.21 s [0.19–0.25]).

### Recording-site localization

We focused on understanding how spiking activity in the primary auditory cortex (A1) contributed to multisensory behavior in behaving primates (269 units; 103 from monkey W and 166 from monkey E). A1 was initially identified based on MRI scans and then functionally by its characteristic anteroposterior tonotopic gradient (Fig. 2A).

**Figure 2:**
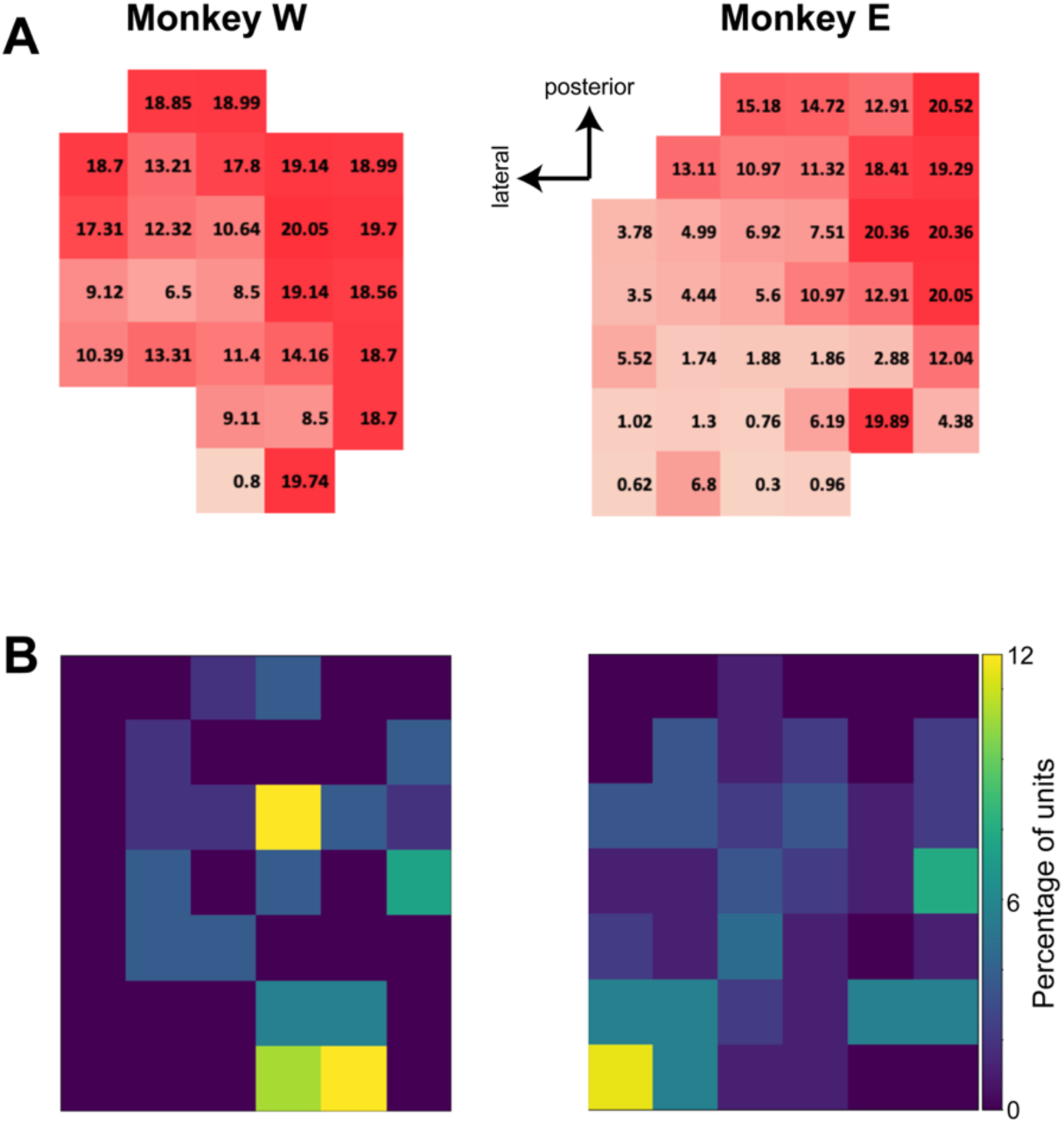
Distribution of best frequency and the proportion of sites with a non-significant STRF in A1. Penetration maps of a monkey’s recording chamber. **(A)** Tonotopic map of the auditory cortex. Each number is the penetration’s best frequency. Color also indicates best frequency: the darker the red, the higher the mean best frequency. **(B)** Percentage of neurons with a non-significant STRF as a function of recording location; see color bar.

We also identified A1 through its current-source-density (CSD) profile (Fig. 3)^18,19^. The A1 CSD profile displayed a dipole pattern indicating net current influx (a putative current sink) and efflux (a putative current source). In the middle channels of the electrode array, the CSD had a sharp negative deflection (current sink) soon after stimulus onset and large increases in MUA (Fig. 3). This large initial current sink and the concomitant large MUA increases are characteristic features of stimulus-evoked laminar response profiles in the core auditory cortex and is consistent with post-synaptic depolarization of neural populations within putative input (granular) layers of the core auditory cortex (lamina 4 and lower lamina 3)^18–22^. Together, these data indicate that we successfully targeted A1 and could align our electrode across the different cortical A1 lamina.

**Figure 3:**
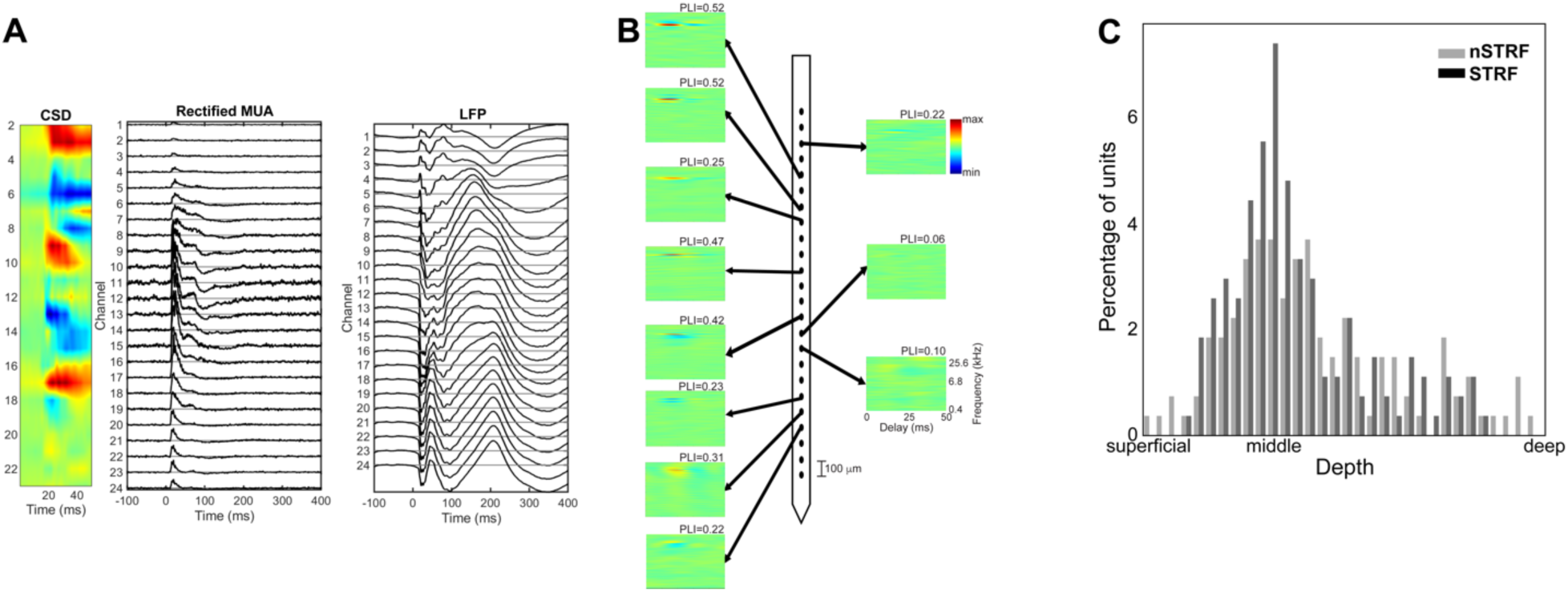
Recording location and distribution of significant and non-significant spectrotemporal response fields (nSTRFs) from a single recording session. **(A)** Representative laminar response profile in A1: **(left)** the two-dimensional current source density (CSD; sinks are color coded in red, and sources are color coded in blue), **(middle)** rectified multiunit activity (MUA), and **(right)** auditory-evoked potentials from the local-field potential (LFP) are shown as a function of channel number from superficial (top) to deep (bottom). The spacing between the electrode channels is 100 µm. **(B)** STRFs and nSTRFs from a single recording session. The panels on the left are significant STRFs, and those on the right are nSTRFs. Arrows indicate channel locations of the recordings on the electrode. The x-axis is aligned relative to stimulus onset. Color indicates the probability of eliciting a spike at each time-frequency point. Each STRF and nSTRF is normalized to its greatest response (red: maximum probability of eliciting a spike; blue: minimum probability of eliciting a spike; see color bar). At the top left of each STRF/nSTRF, we provide its phase-locking index (PLI). The scale bar indicates the 100-µm spacing between electrode channels. **(C)** Distribution of nSTRFs and STRFs across all recordings and both monkeys as a function of cortical depth. Each session was aligned to the maximum MUA and the earliest source CSD (see Fig. 3). Grey (black) data are nSTRFs (STRFs).

### Neurons with STRFs and non-significant STRFs (nSTRFs) originate from distinct neural A1 populations

Because frequency-response fields are a fundamental organizing principle of the auditory system, it is important to understand their relationship with behavior^23^. To this end, in each recording session, we delivered dynamic moving ripple (DMR) stimuli and constructed each neuron’s STRF^24,25^ from the recorded activity. Although such measurements are commonplace in other studies and species, the nature of STRFs in the primate A1 is not well described.

We used a two-step procedure to identify the reliable samples (pixels) for each STRF and its overall reliability^25–27^; more details of this procedure can be found in **Materials and Methods**. In brief, a bootstrap resampling procedure identified STRF samples that were significant and in a second step, we accounted for false positives using a reliability analysis. An example recording session is shown in Fig. 3B. This figure shows the STRFs and non-significant STRFs (nSTRFs) from all of the simultaneously recorded single units. We found that nSTRFs were relatively common: on average, ∼47% of single units had nSTRFs. We identified STRFs and nSTRFs across all the cortical layers and at all best frequencies (Fig. 2B, C). We could not identify any difference between the laminar distribution of STRFs and those of nSTRFs (χ^2^-test, H_0_: the proportion of STRFs and nSTRFs as a function of depth is the same; p>0.05; Fig. 3C).

A basic requirement underlying the construction of a significant STRF is that a neuron’s action potentials time lock (phase lock) precisely to the envelope of the DMR stimulus. To test the hypothesis that the difference between our STRF and nSTRF populations can be, at least partially, attributed to this mechanistic difference, we computed a phase-locking index for each STRF and nSTRF^25^. A phase-locking index value of 0 indicates that there was a poor relationship between the timing of action potentials and instances of the DMR envelope, whereas a value of 1 indicates a tight relationship between action-potential timing and the DMR envelope. Consistent with our hypothesis, we found that the median [IQR] index value for STRF neurons (0.274 [0.16–0.37]) was significantly higher (Mann-Whitney test, H_0_: median phase-locking index values were the same, p<0.0001) than the median index value for nSTRF neurons (0.074 [0.05–0.10]).

We conducted three additional analyses to explore differences between the STRF and nSTRF neurons. First, we found that the firing rate of STRF neurons during the DMR stimulus (median [IQR]: 9.52 spikes/sec [4.3–15.8]) was significantly higher (Mann-Whitney test, H_0_: median firing rates for STRF and nSTRFs were the same; p<0.000002) than the firing rate of nSTRF neurons (median [IQR]: 4.8 spikes/sec [2.3–9.6]). In the second analysis, we tested whether STRF and nSTRF neurons have distinct action-potential waveforms, which could indicate that these neural populations originate from distinct neural populations^28–30^. Indeed, as shown in Fig. 4, we found that the median [IQR] trough-to-peak duration of nSTRF neurons (0.45 ms [0.20–0.70]) was longer (Mann-Whitney test; H_0_: median durations were the same; p<0.0001) than that of STRF neurons (median [IQR] = 0.21 ms [0.16–0.37]). Finally, we asked whether STRF and nSTRF neurons had different functional organizations^31^. To assess differences in functional organization that may relate to different patterns of anatomical connnectivity^32^, we measured the noise correlation between simultaneously recorded pairs of neurons (Fig. 5). For STRF neurons during difficult trials (i.e., SNR: -15 and -10 dB), we could not identify a difference between the median noise correlation during visually congruent trials from that during visual-static trials (Wilcoxon test; H_0_: median noise correlations were the same; p>0.05 for both monkeys and individually for monkey W and monkey E). In contrast, for nSTRF neurons during low SNR (-10 and -15 dB; i.e., difficult) trials, the median noise correlation during visually congruent trials was higher than during visual-static trials (Wilcoxon test, p<0.001 across both monkeys; Monkey W: p<0.003; Monkey E: p<0.04). We could not identify differences in noise-correlation values during higher SNR (i.e., easier) trials, consistent with previous work demonstrating that noise correlations are highly dependent on stimulus-task interactions^33,34^.

**Figure 4:**
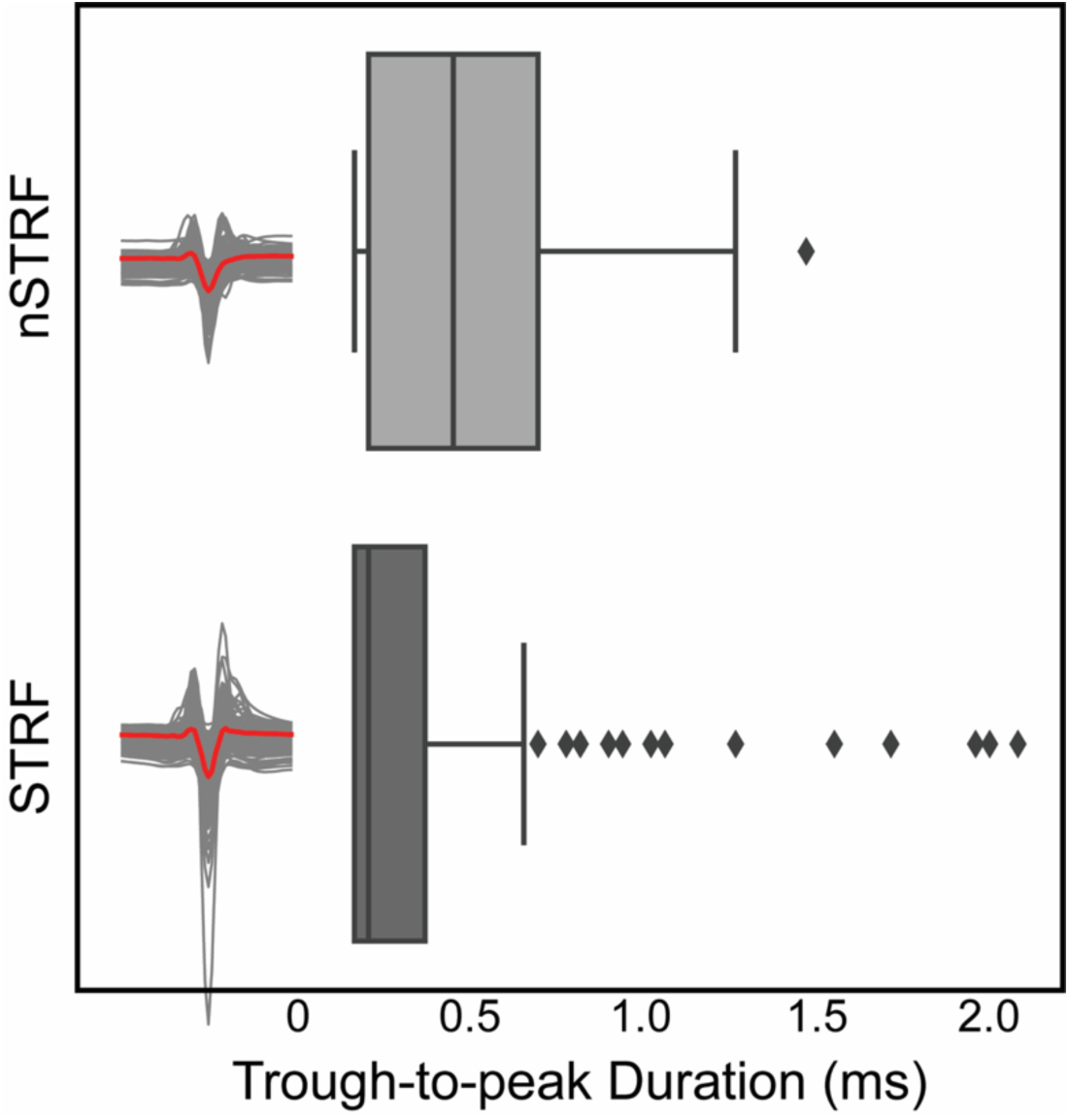
nSTRFs and STRFs neurons have different spike waveforms. Individual spike waveforms (black) and the median spike waveforms (red) of nSTRF and STRF A1 units. A whisker plot illustrates that the median trough-to-peak duration of nSTRF neurons is longer (Mann-Whitney test, p<0.0001) than those with STRFs.

**Figure 5:**
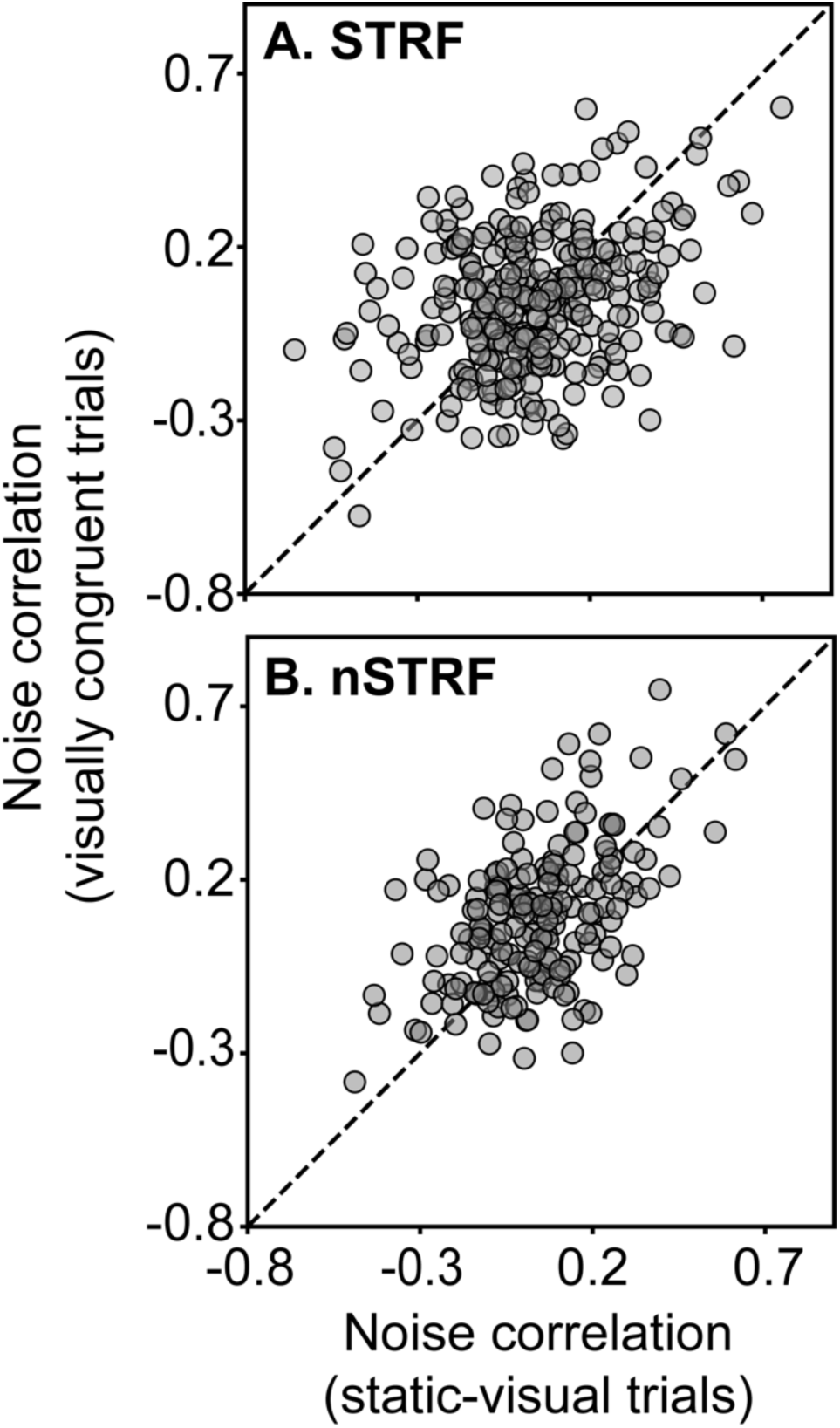
nSTRF and STRF neurons have different patterns of functional connectivity. **(A)** A scatterplot showing the relationship between the noise correlation of simultaneously recorded pairs of STRF neurons during static-visual trials (median [IQR] = 0.15 [-0.09–0.17]; x-axis) and during visually congruent trials (0.047 [-0.08–0.19]; y-axis) for difficult SNR trials. **(B)** Same as **A** but for nSTRF neurons (static-visual trials: 0.036 [-0.08–0.15]; visually congruent trials: 0.1 [-0.06–0.21]). In both plots, each data point represents a single pair of simultaneously recorded neurons; data are collapsed across both monkeys and all sessions. The dotted line is the line of equality.

In subsequent analyses, we evaluated STRF and nSTRF A1 neurons independently to test whether they differentially contribute to multisensory behavior.

### The spiking activity of nSTRF neurons is modulated more by visual-trial condition (visually congruent versus static-visual trials) than STRF neurons

We discovered that the multimodal task altered STRF and nSTRF neurons in different ways. Two example neurons are shown in Fig. 6. Both example neurons showed robust time-locked responses to the onset of the chorus (Fig. 6, left). Later in the task, when we presented the vocalization target, we found fewer time-locked responses but clear modulation by SNR (Fig. 6, middle and right). The biggest difference between these STRF and nSTRF example neurons was the degree of modulation by the visual stimulus in the static-visual trial (Fig. 6, middle) and congruently visual trial (Fig. 6, right): the nSTRF neuron was modulated more during the congruently visual trial, especially at the lower (i.e., more difficult) SNR values.

**Figure 6:**
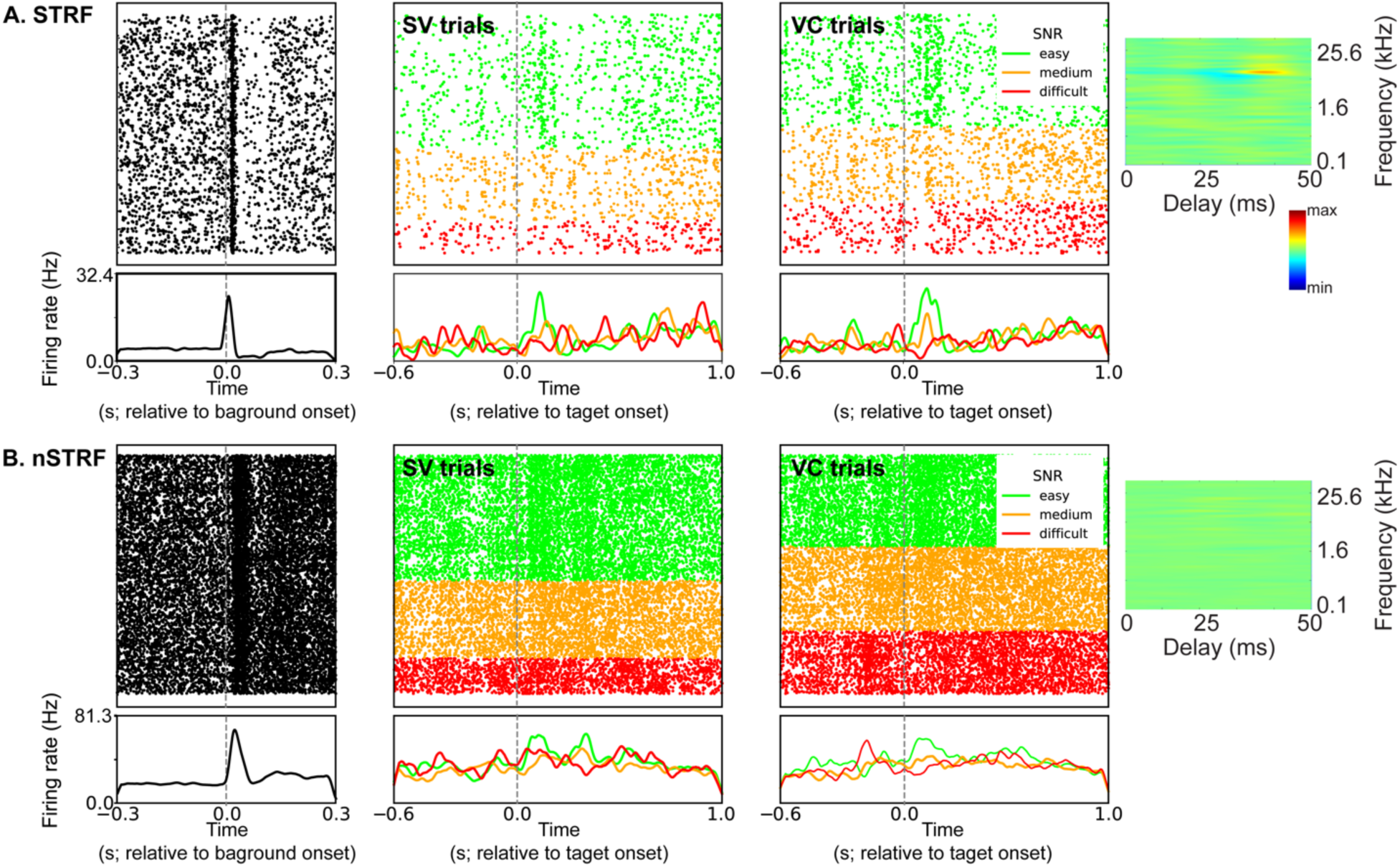
Single-neuron examples. **(left)** Raster and peristimulus time histogram (PSTH) relative to onset of the background chorus. **(middle)** Rasters and PSTHs from the static-visual (SV) trials. The rasters and PSTHs are organized as a function trial difficulty: “easy” (SNR= 5 and 10 dB), “medium” (SNR= -5 and 0 dB), and “difficult” (SNR= -15 and -10 dB) trials. **(right)** Rasters and PSTH from the visually congruent (VC) trials. Same organization as the **middle** panel. At the **far right**, we have plotted each neuron’s **(top)** STRF and **(bottom)** nSTRF. The x-axes for both are aligned relative to stimulus onset. Color indicates the probability of eliciting a spike at each time-frequency point. Each is normalized to its greatest response (red: maximum probability of eliciting a spike; blue: minimum probability of eliciting a spike; see color bar). All data are from correct trials only.

To quantify this observation across our population of neurons, we calculated a “modulation index” (Fig. 7). This index quantified the degree to which firing rate was modulated by congruently visual trials versus the static-visual trials. An index value of 0 means that a neuron was modulated similarly by two visual trial conditions. Larger positive (negative) values mean that a neuron was modulated more during visually congruent (static-visual) trials. A linear-mixed effects model (SNR value and neuron class [STRF versus nSTRF] as main effects with number of single units in each group as random effects) indicated that the median modulation-index value increased with SNR (H_0_: median index value is the same across SNR levels; p<0.004) and that the median modulation-index value for nSTRFs was larger than for STRF neurons (H_0_: median index value was the same for STRF and nSTRF neurons; p<0.02). We could not identify a robust relationship between cortical (laminar) depth and modulation-index value (data not shown).

**Figure 7:**
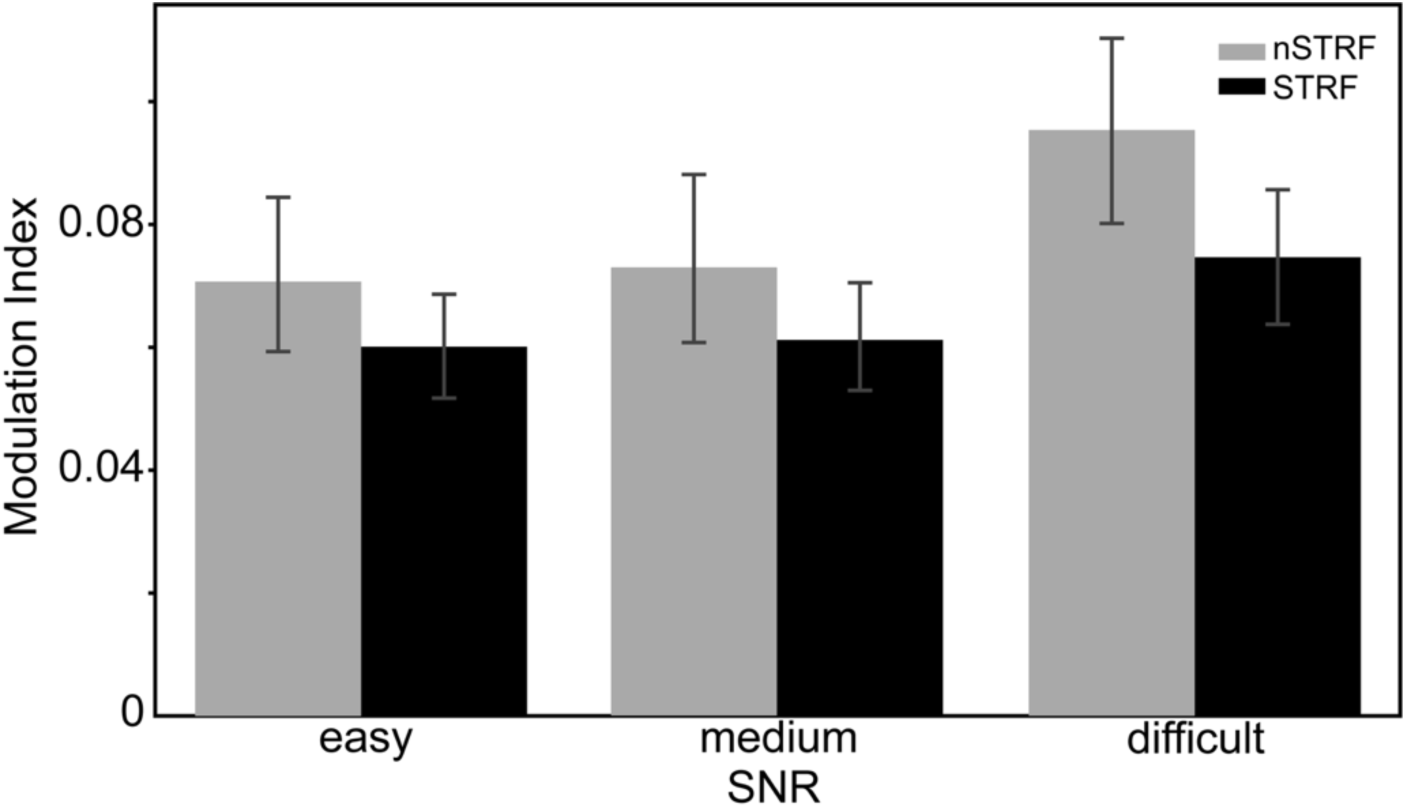
nSTRF neurons have larger modulation-index values than STRF neurons. Bar plots of modulation-index values as a function of “easy” (SNR= 5 and 10 dB), “medium” (SNR= -5 and 0 dB), and “difficult” (SNR= -15 and -10 dB) trials. Error bars indicate 95% bootstrapped confidence interval. Easy-trial modulation-index values (median [IQR]): STRF = 0.05 [0.02–0.09] and nSTRF = 0.05 [0.02–0.09]. Medium-trial modulation-index values (median [IQR]): STRF = 0.05 [0.02–0.08] and nSTRF = 0.04 [0.02–0.09]. Difficult-trial modulation-index values (median [IQR]): STRF = 0.05 [0.02–0.11] and nSTRF = 0.06 [0.03–0.13].

### The population trajectories of STRF and nSTRF neurons encode SNR value comparably. However, the trajectories of nSTRF neurons are more robustly modulated by the different visual trial conditions than the STRF-neuron trajectories

A powerful approach to understanding how information is encoded by neural populations is to train classifiers (support-vector machines) to decode a particular task variable (for example, different SNR values). However, one of drawbacks of this approach is that we are unsure of how much of the trained decoding space represents the variation in neural activity brought about by the task variable of interest (e.g., SNR value) as opposed to the variation in neural activity brought about by the interaction of the other variables (i.e., choice, visual-trial condition, etc.) with the trained decoding space.

To overcome this limitation, we conducted an analysis in which we represented the firing rates of N recorded neurons as a time-varying trajectory in a state space that is spanned by orthogonal axes. These orthogonal axes are defined by the task variables^35^. In brief, we first generated a lower-dimensional representation of the space. This space was then projected onto the orthogonal axes that were defined by the variables (e.g., SNR, visual-trial condition, choice) of our multisensory task. As a consequence, we could isolate the representation of one task variable, independent of the other task variables.

The results of this analysis are shown in Fig. 8. The trajectories of both STRF and nSTRF neural populations were modulated by the SNR between the target and the background chorus (bottom panels of Figs. 8A and 8B). That is, over time, the distance between the neural trajectories and the -15-dB trajectory increased. The amount of separation is a function of SNR: as the SNR value increased, the trajectory separation increased. The separation peaked a few hundred milliseconds after onset of the target vocalization and then returned to baseline (i.e. no separation).

**Figure 8:**
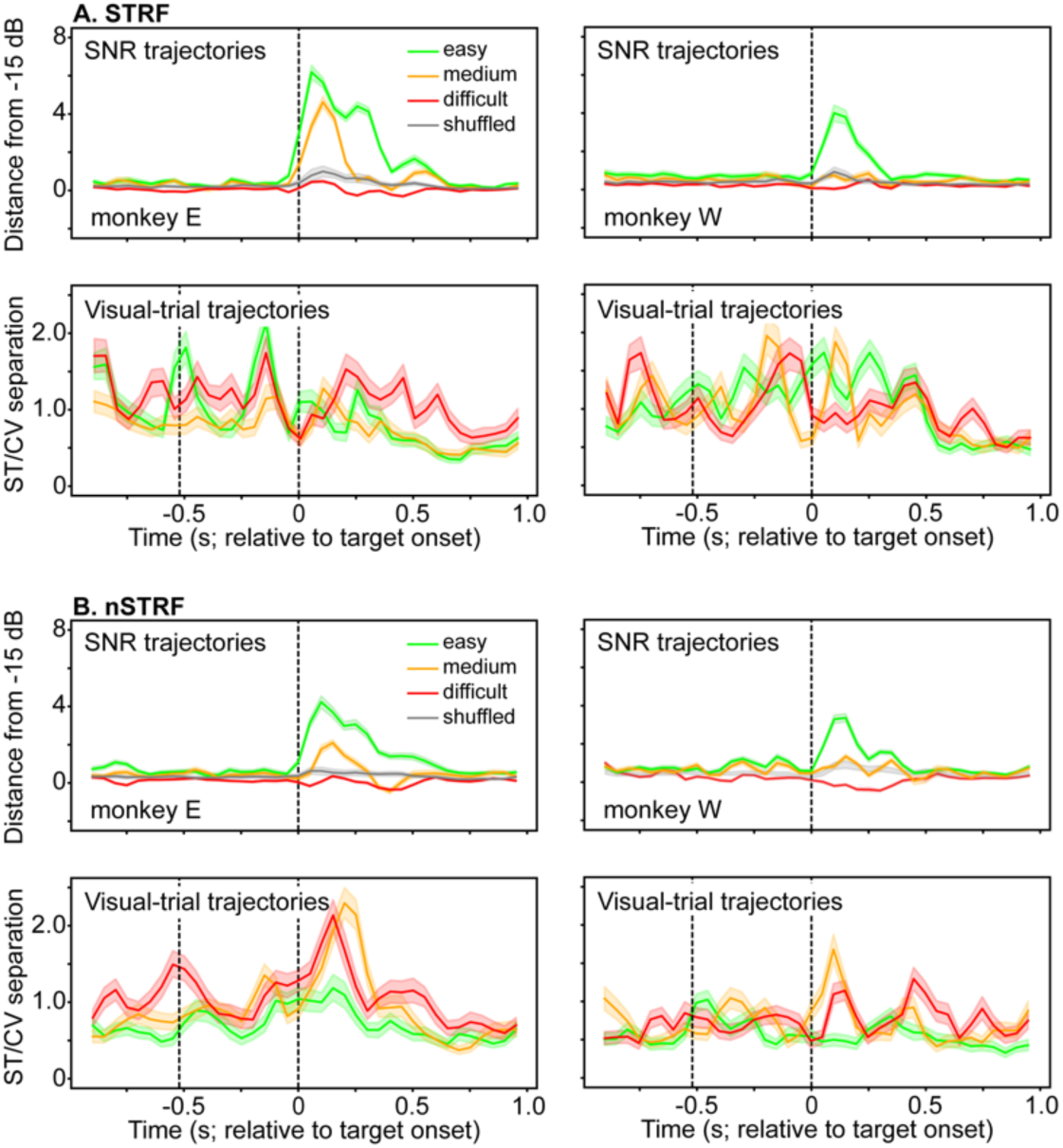
Population trajectories of STRF and nSTRF neurons encode SNR value comparably. *However*, the trajectories of nSTRF neurons are modulated more by the visual stimulus than the STRF-neuron trajectories. (A, top panels: SNR trajectories) The average distance of the population STRF trajectories for “easy” (SNR= 5 and 10 dB), “medium” (SNR= -5 and 0 dB), and “difficult” (SNR= -15 and -10 dB) trials, relative to the -15 dB SNR trials. The solid lines represent mean values, whereas shading indicates the 95% bootstrapped confidence intervals. Grey indicates the average and 95% confidence interval for a bootstrapped null distribution. The vertical dotted line indicates onset of the target coo. **(A, bottom panels: Visual-trial trajectories)** The average distance between the population trajectories during visually congruent (VC) and static visual (SV) trials as a function of easy, medium, and difficult trials. The solid lines are mean values, whereas shading indicates the 95% bootstrapped confidence intervals. The first vertical dotted line indicates the onset of the video and the second indicates the onset of the target coo. Data in **B** are organized the same as in **A** but for nSTRF neurons. All trajectories are aligned relative to onset of the target coo.

However, a different picture emerged when we examined the STRF and nSTRF trajectories of the static-visual and visually congruent trial conditions (bottom panels in Fig. 8A and Fig. 8B). Here, we see that the nSTRF trajectories were modulated more by the different visual-trial conditions than the STRF trajectories. Moreover, although these trajectories had some local peaks, the largest peak in the nSTRF trajectories (but not the STRF trajectories) occurred after target onset. This peak was also temporally aligned with the peak of the SNR trajectories. Indeed, we found that the peak of the SNR trajectories and the peak of the visual-trial trajectories were more temporally aligned for the nSTRF trajectories than for the STRF trajectories (Mann-Whitney test; H_0_: time difference between the SNR and visual-trial trajectories is the same for STRF and nSTRF trajectories; easy SNR-trial trajectory difference; Monkey E: p<1.9×10^-11^, Monkey W: p<0.03; medium SNR-trial trajectory difference; Monkey E: p<0.000002, Monkey W: p <0.02; difficult SNR-trial trajectory difference; Monkeys E and W: p>0.05). We note that for the difficult trials, this analysis was compromised because there was not a peak for the difficult SNR-trial trajectories (see top panels in Fig. 8).

The nSTRF visual trajectories were also systematically modulated by the SNR value. As SNR values decreased, there was a larger separation between the trajectories of the two visual-trial conditions. In other words, as the target vocalization became harder to detect (i.e., with decreasing SNR values), there was a concomitant increase in the representation of visual information in the trajectories. Together, this suggests a correspondence between these two variables with respect to the multisensory task.

Finally, we asked whether the trajectories differed as a function of the monkeys’ choices (Fig. 9). We found, for both STRF and nSTRF neurons, that neural trajectories on hit trials were distinct from those on miss trials. However, this separation did not occur until after the decision time (the vertical dotted line in Fig. 9).

**Figure 9:**
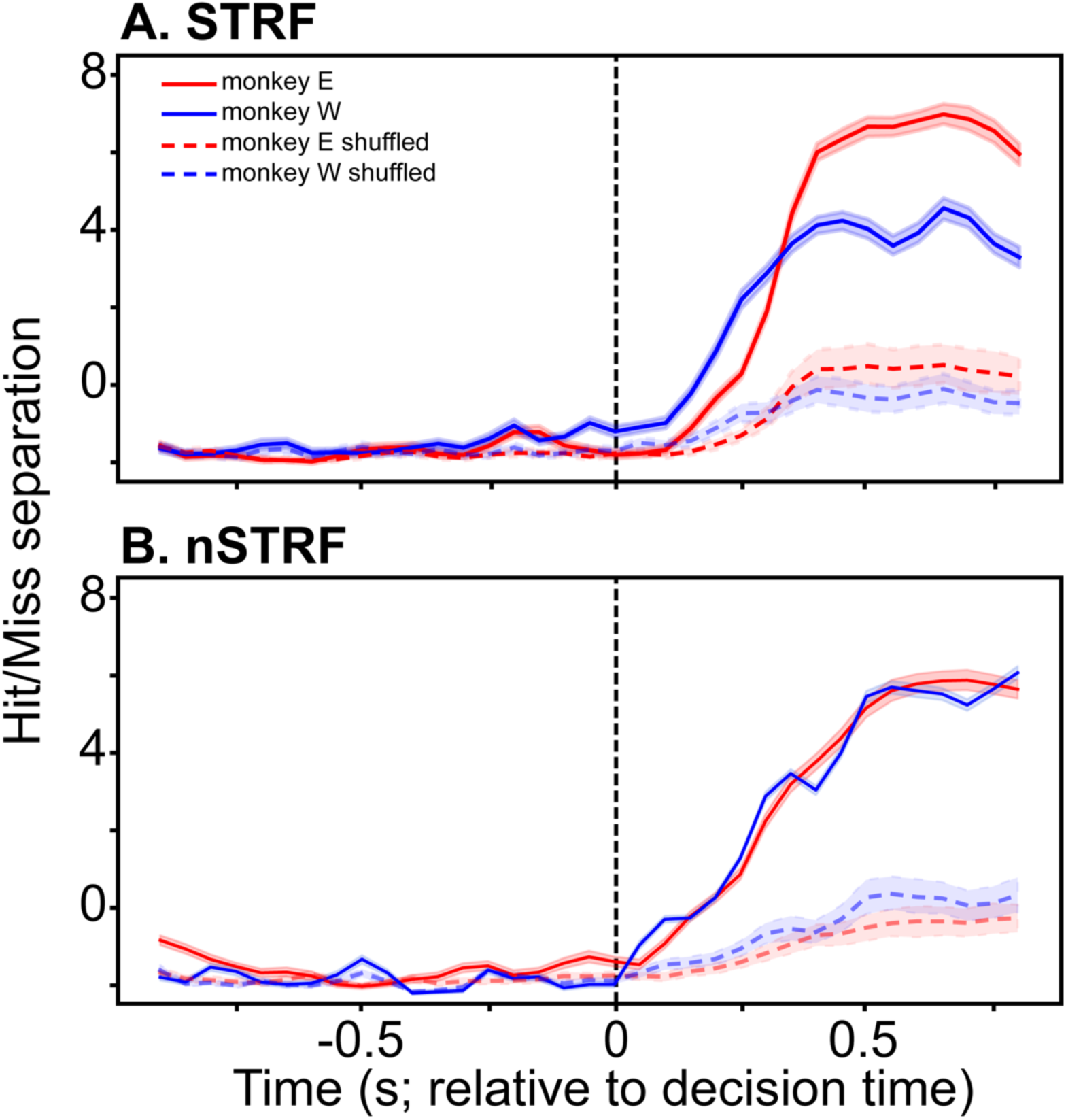
The population trajectories of STRF and nSTRF neurons encode choice after the decision time. **(A)** The average distance between the population trajectories during hit and miss trials. The solid lines are mean values, whereas shading indicates the 95% bootstrapped confidence intervals. The dotted lines and shading indicate the mean and 95% confidence interval for a bootstrapped null distribution. Data in **B** are organized the same as in **A** but for nSTRF neurons. All trajectories are aligned relative to the decision time (i.e., the vertical dotted line).

## Discussion

We combined behavior and neural recordings to test how A1 spiking activity contributes to an ethologically inspired multisensory detection (source-segregation) task. We found that the monkeys’ hit rates were higher during congruently visual trials than during static-visual trials. This was most apparent at low SNR values, indicating that, like other multisensory studies, the effects of multisensory stimuli on behavior are most prominent when a unimodal stimulus is weak^2^. The benefit of the congruent (dynamic) video is also consistent with a large body of research showing that congruent mouth movements facilitate speech/vocalization perception^3,36–44^.

In addition to higher hit rates, we found that the monkeys’ decision times, which in a drift-diffusion model is the time between stimulus onset and the time to cross the decision bound^14,15^, were faster during congruently visual trials than during static-visual trials. To our knowledge, this is the first application of a single-choice drift-diffusion model to multisensory processing. This result is consistent with previous work showing that, relative to unimodal decisions, subjects respond faster during multisensory decisions^45,46^. It is possible that these faster response times may also be attributed to changes in decision times.

We can compare the decision time with the temporal dynamics of A1 activity to gain insights into its contribution to our multisensory perceptual decision^47^. Because differences between the hit and miss trajectories were only apparent after the decision time (see Fig. 9), A1 activity does not appear to reflect a component of a feedforward decision process. Instead, a more parsimonious explanation is that these signals represent feedback to A1 after the decision is formed elsewhere^48,49^. In contrast, our previous work indicated that choice-related modulations in the anterolateral belt region of the auditory cortex occurred near or just before the decision time, suggestive of a feedforward contribution to the decision^50^. Additional work is needed to identify the brain region(s) that form our multisensory decision and to identify the neural mechanisms and computations that underlie this decision.

In auditory neurophysiological studies, it is a common strategy to restrict analyses to neurons that have a clear auditory response. In some studies, investigators may first construct a neuron’s linear frequency response field and then compute the neuron’s best frequency from the response field. If there is a best frequency, the investigator might then titrate a subsequent stimulus relative to this best frequency. For example, if a neuron’s best frequency is 1.5 kHz, the target stimulus in a behavioral task may be a 1.5-kHz tone burst. Other studies may not construct the response fields but only analyze neurons that respond to the target stimulus^51^.

Inherently, this strategy limits analyses to only the subpopulation of neurons that are “auditory”^51^. Although this strategy can be powerful, we chose not to follow it for two reasons. First, our goal was to study an ethologically relevant sound-segregation scenario that utilizes species-specific vocalizations during naturally occurring multisensory integration. Since both the target and chorus stimuli for this task were broadband, it did not make sense to titrate our stimuli to the A1 neurons’ best frequency. Second, because we found that a large proportion of our neurons did not have significant STRFs, which to our knowledge has not been documented, we were curious to determine whether and how they contributed to behavior. Indeed, a neuron’s response properties during passive listening (which occurred when the monkeys listened to the DMR stimulus) may be quite different from those during active listening (i.e., during their participation in the multisensory task)^52–56^.

We found that STRF and nSTRF neurons differed considerably with respect to their contribution to multisensory behavior. In particular, individual and populations of nSTRF neurons were modulated more by the different visual-trial types (visual static versus congruently visual) than STRF neurons. This was most evident in the visual-trial nSTRF trajectories, where we identified modulatory increases as a function of the SNR between the target vocalization and the background chorus. (Fig. 8). These increases in the nSTRF trajectories (but not the STRF trajectories) occurred after target onset and at approximately the same time as the peak of the SNR trajectories; compare Figs. 8A and 8B.

Do STRF and nSTRF neurons originate from the same class of neuron? Or do they represent distinct neural populations? We suggest that our data are more consistent with the latter proposition because (1) they had different firing rates, (2) the shapes of the STRF and nSTRF spike waveforms were different, and (3) they had different patterns of functional connectivity (see Figs. 4 and 5). This functional distinction between neural populations is akin to our previous work showing that putative auditory-cortex interneurons are more category selective than putative pyramidal neurons^30^. This nSTRF-STRF proposition adds to our understanding of the functional organization of cortical microcircuits in the auditory cortex: whereas previous work has examined how spiking activity in cortical microcircuits was modulated by a stimulus’ features^57–59^, we posit that this functional differentiation between STRF and nSTRF neurons sheds light on how different elements of the A1 cortical column may contribute to auditory/multisensory behavior.

Of course, we cannot definitively know whether nSTRF and STRF neurons belong to distinct populations and would need viral interrogation or other probes to positively identify whether they belong to distinct neural classes. It is also possible that we might have identified response fields in the nSTRF neurons if we had presented a different stimulus set such as tone bursts or dynamic random chords^25,60,61^. However, even this possibility would still indicate some fundamentally different type and possibly very nonlinear type of integration that differentiates STRFs and nSTRF neurons. Finally, we cannot attribute this finding to simple sampling bias: we identified STRF and nSTRF neurons during the same recording session and found nSTRFs throughout all the cortical layers (see Figs. 2 and 3).

Overall, our findings are the first to demonstrate how functional information relating to multisensory behavior is organized in the primate A1, to identify a differential contribution of nSTRFs to behavior, and to demonstrate that task-related information in the primate A1 is encoded as a structured dynamic process in the neural population. Further work is needed to determine whether there is also a differential causal role in multisensory behavior between STRF and nSTRF neurons, whether nSTRF neurons are, in general, more susceptible to extra-sensory modulation^62–64^, the degree to which a neuron’s linear response field (e.g., a STRF) relates to its contribution to behavior, and if STRF neurons reflect distinct stages of processing within the circuitry of the A1 cortical column.

## Materials and Methods

The University of Pennsylvania Institutional Animal Care and Use Committee approved all the experimental protocols. All surgical procedures were conducted under general anesthesia and using aseptic surgical techniques. The authors were not blind to group allocation during the experiment and when analyzing the data outcomes.

### Experimental chamber

In each session, a male monkey (*Macaca mulatta*; monkey W [9 years old] or monkey E [9 years old]) was seated in a primate chair. A Dell monitor (DELL E1709W, 59.89 Hz) and a calibrated free-field speaker (MICRO [Gallo Acoustics]) were placed 1 m in front of the monkey and at their eye level. The monkey moved a joystick, which was attached to the primate chair, to indicate their behavioral report. These sessions took place in an RF-shielded room that had sound-attenuating walls and echo-absorbing foam on the inner walls.

Visual stimuli were delivered through customized python (3.6.13) scripts and Psychopy (2021.1.4), which were ultimately presented on the Dell monitor. Calibrated auditory stimuli were generated with Matlab (Mathworks), passed through a digital-to-analog convertor (E22, Lynx Studio Technology Inc), and presented through the free-field speaker.

### Identification of A1

We identified the stereotactic location of the core auditory cortex (A1) through MRI scans^65–69^. We functionally defined A1 by a variety of metrics. First, A1 neurons have short-latency responses and circumscribed spectrotemporal response fields (Fig. 3B). Second, using data from different recording sessions, we generated A1’s characteristic anteroposterior tonotopic gradient (Fig. 2A). Third, because we oriented our electrode orthogonal to A1’s cortical layers (lamina), A1 could be identified by its laminar response profile, including its current-source density (CSD; which is the second spatial derivative, relative to electrode-contact separation, of the local-field potential) as well as the spatial pattern of multiunit activity (Fig. 3A)^18,19,21,22,70^.

### Recording methodology and strategy

During a recording session, we penetrated the dura with a stainless-steel guide tube (508-µm inner diameter) and then advanced the electrode (24-channel v-probe, 100-µm equal spacing between channel [Plexon]) inside the guide tube. The neural signals were amplified (PZ5, Tucker-Davis Technologies), digitized, and stored (sampling rate: 24.4 kHz; RZ2, Tucker-Davis Technologies) for online and offline analyses.

While advancing the electrode, we presented an auditory ‘‘search stimulus’’ (80-ms white noise burst; 65 dB SPL). Once we found auditory responses, we generated the site’s current-source density (via ∼130 repetitions of the search stimulus) and analyzed the resultant CSD and multiunit activity. We adjusted the v-probe depth to help to ensure that the highest rate of multiunit activity and the initial short-latency CSD sink were aligned with the middle channels of the v-probe. This initial short-latency current sink represents extracellular current flow associated with post-synaptic potentials in stellate and pyramidal neurons with cell bodies located in the thalamorecipient zone (i.e., lamina 4 and lower lamina)^18,19^. Once we identified this location, we retracted the electrode by 250-500 µm and allowed the tissue to stabilize for >50 mins to minimize electrode drifting.

We then collected neural activity while the monkey listened passively to an auditory stimulus. We used this neural activity to construct the spectrotemporal response fields from the spiking activity of isolated single neurons. Next, the monkey participated in trials of our multisensory task. At the end of a recording session, we, once again, presented the auditory stimulus to reconstruct the spectrotemporal response fields and regenerated the CSD and multiunit profiles.

### Dynamic moving ripple (DMR) noise: stimulus to generate a neuron’s spectrotemporal response field (STRF)

As we just mentioned, while a monkey listened passively, we presented a DMR noise stimulus and collected spiking activity from single units^24,25^. We used this spiking activity to construct a neuron’s STRF.

DMR noise is a continuous time-varying broadband noise stimulus that covers the frequency range between 0.1 and 32 kHz (10-min duration; 65 dB spectrum level per ⅓ octave; 96-kHz sampling rate; 24-bit resolution)^25,27^. At any instant of time, the stimulus had a sinusoidal spectrum; the density of the spectral peaks was determined by the spectral modulation frequency (0-4 cycles/octave). The peak-to-peak amplitude of the ripple envelope was 30 dB. The stimulus also contained temporal modulations that were controlled by the temporal modulation frequency (0-50 Hz). Both the spectral and temporal parameters were unbiased (uniformly distributed) and varied randomly and dynamically over time; the maximum rates of change were 0.25 Hz and 1 Hz, respectively.

### Multisensory task: stimuli and task structure

In our multisensory task (Fig. 1A), monkeys reported the presence of a “target” vocalization that was embedded in a “background chorus” of vocalizations. The monkey moved the joystick to indicate their detection of the target vocalization. We trained the monkeys to respond as fast as possible; i.e., this was a reaction-time task.

On a trial-by-trial basis, we randomly varied the sound level of the target vocalization, relative to the level of the background chorus (65 dB SPL). The signal-to-noise ratio (SNR) between the target and the background was between -15 and 10 dB, with a step size of 5 dB. We paired the target-background stimulus with different visual stimuli, which also titrated task difficulty. This also randomly varied on a trial-by-trial basis.

#### Auditory stimuli: target vocalization and background chorus

The target vocalization was a single exemplar of a *coo* (duration: 700 ms) that was obtained from a video recording of an unknown conspecific^71^. The background chorus was created by randomly superimposing 30-40 different vocalizations, which were also from unknown conspecifics^72^. We minimized the amplitude troughs of this stimulus mixture to reduce the possibility that a monkey could detect the target vocalization if it occurred within an amplitude trough of the background chorus (i.e., glimpsing)^73–75^. We created new tokens of the background chorus for each stimulus condition.

#### Trial conditions

There were 3 different trial conditions based on the relationship between the target coo vocalization and the accompanying visual stimulus. Each trial started with the background chorus and a static visual image of the monkey’s face with a closed mouth.

During “static-visual” trials, this static image of the monkey’s face with a closed mouth remained on throughout the trial. We presented the target coo at a variable delay of 1-2.5s after onset of the background chorus.

During “visually congruent” trials, the target coo was accompanied by a dynamic video of the monkey moving its mouth synchronously as it vocalized the target coo. Like the “static visual” condition, we presented the target coo and the congruent video of the monkey vocalizing the coo at a variable delay of 1-2.5 s after chorus onset.

During “visual-only” trials (∼20% per session), we presented the dynamic video of the monkey cooing <u>without</u> the accompanying target coo (i.e., auditory stimulus). 500 ms after the offset of the video, we presented the coo (SNR = 10 dB only). This was a *catch* trial to test for the possibility that the monkeys responded to the dynamics of the video and not to the target coo itself.

#### Trial outcomes

The multisensory task was a “yes/no” detection task. During visually congruent and static-visual trials, a “hit” occurred when we presented the target coo vocalization and the monkey moved the joystick (i.e., a “yes” report) within 800 ms of target onset. In contrast, on these trials, a “miss” occurred when we presented the target vocalization, but the monkey did not move the joystick within 800 ms. During visual-only trials, a “correct rejection” occurred when the monkey maintained their grip on the joystick and moved it only in response to the coo at the end of the trial. On any trial, a “false alarm” occurred when the monkey moved the joystick before presentation of the coo. We rewarded monkeys on hit and correct-rejection trials. On false-alarm and miss trials, we added an extra 2 s to the inter-trial interval.

### Behavioral analysis

#### Psychometric data

We fit the behavioral data to a Weibull cumulative distribution that describes the relationship between hit rate (i.e., number of hits divided by the number of hits and misses) and SNR^76^: 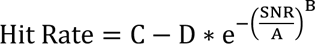. C and D represent the lower and upper asymptotes, respectively. A relates to the horizontal position of the curve, and B modifies the shape of the function. We independently fit session-by-session data from the static-visual trials and visually congruent trials.

#### Decision time

We analyzed behavior using a variant of a sequential-sampling model of decision-making^77–80^. Sequential-sampling models quantify the process of converting incoming sensory evidence, which is represented in the brain as the noisy spiking activity of sensory neurons, into a decision variable that guides behavior. Here, we used a standard drift-diffusion model, in which the decision is based on a temporal accumulation of noisy evidence to a fixed bound. This process is mathematically equivalent to the one-dimensional movement of a particle undergoing Brownian motion to a boundary. We utilized a model version that incorporates a single choice (i.e., the detection of the target vocalization)^14,15^. “Decision time” was defined as the time between stimulus onset (which in our case is the target vocalization) and the time to cross the bound (i.e., the decision criterion). “Non-decision time” reflects sensory encoding, sensorimotor transformations, and other components that are unrelated to the decision. “Response time” was the sum of the decision time and the non-decision time. We assumed that the rate of evidence accumulation (drift rate) varied across trials and followed a normal distribution N(ξ,η), where ξ is the mean drift rate and η is the standard deviation. We also assumed that the non-decision time varied across trials according to a uniform distribution 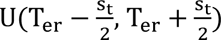, where T_er_ is the mean and s_(_ is the width of the distribution. In each session, as a function of trial condition, we fit this DDM model to the response-time distributions with a fixed-value decision criterion.

### Neurophysiological data analysis

Neural signals were zero-phase filtered between 300-5000 Hz with a 20^th^-order FIR filter and then sorted offline with Kilosort3^81^. We manually inspected all sorted units for stability over the duration of the recording session. We did not use statistical methods to predetermine sample sizes, but our sample sizes are like those reported in previous publications and are similar to those generally employed in the field^50,82–84^.

#### Method to generate a STRF from single-unit spiking activity

For each neuron, we used reverse correlation to construct its STRF, which is the average spectrotemporal stimulus envelope immediately preceding each spike^25,26^. We utilized a two-step procedure to identify each STRF’s reliable samples and its overall reliability^27^. To identify the significant spectrotemporal samples in each STRF, we generated a null model under the assumption that spiking activity occurred randomly with respect to the DMR. We calculated this null model by randomizing the inter-spike intervals and then averaging the DMR’s spectrotemporal envelope relative to the time of each spike. This generated a “noise” STRF. We repeated this procedure 1000 times to generate a distribution of spectrotemporal noise-STRF samples. We considered a STRF sample significant if its value was outside of the 99.9% confidence interval (p<0.001) of its respective noise-STRF sample. STRF samples that overlapped with the noise-STRF distribution were assigned a value of 0.

For any given STRF, ∼16 STRF samples (i.e., 200 temporal × 83 spectral samples × 0.001) were false positives; that is, they exceeded the significance level of p<0.001 by chance. Although these samples exceeded our significance criterion, they did not fulfill our primary objective of identifying STRFs with reproducible and reliable auditory responses. To identify such STRFs, we computed a “reliability” index to identify a STRF with reproducible structure. We calculated this index by breaking the ripple stimulus into twenty 30-s long segments (10-min total). We next randomly selected two subsamples of the 20 segments (subsample A and B, each containing 10 DMR segments chosen randomly), generated a STRF from each of these segments (STRF_A_ and STRF_B_), and calculated the Pearson correlation coefficient between STRF_A_ and STRF_B_. A correlation coefficient ∼1 indicates that a STRF was very reliable, consistent with a stable stimulus-evoked and synchronized response, whereas a value ∼0 indicated that it was not reliable. To generate a distribution of correlation-coefficient values, we repeated this procedure 500 times. We next generated a null distribution of coefficient values using the same process but with randomized inter-spike intervals. A Mann-Whitney test evaluated the null hypothesis that the actual distribution of correlation-coefficient values and the null distribution had the same median values. If the null hypothesis was rejected (p<0.01), we considered a STRF to be “reliable”.

#### Single-neuron analysis: task modulation index

On a neuron-by-neuron basis, we calculated a neural modulation index MI: 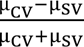, where µ is the mean firing rate over the 300-ms window starting at target-vocalization onset during congruently visual (CV) and static-visual (SV) trials, respectively. We employed a bootstrapped procedure to estimate the MI for each neuron. If the index value ∼0, it implied that the firing rate was not modulated by trial condition, whereas more positive (negative) values indicated that the firing rate during visually congruent trials was greater (less) than during static-visual trials. We used a linear mixed-effects model with SNR and STRF significance (SIG_STRF_) as fixed effects and the number of single units as random effects to evaluate the null hypothesis that the median MI value was the same for STRF and nSTRF neurons and across all SNRs: MI∼SNR + SIG_STRF_ +SNR× SIG_STRF_ +(1|number of single units).

#### Targeted-dimensional reduction of population activity

A neural trajectory is the N-dimensional curve traced by the spiking activity of N neurons as a function of time. We instantiated a dimensionality-reduction technique to find the dimensions in this space that were most informative and to test how these trajectories were modulated by a task variable^35^. Using a bootstrap procedure, we generated 100 pseudopopulations of 30 STRF and 30 nSTRF neurons by resampling across recording sessions.

We first constructed a matrix (**M**) of z-scored firing rates as a function of time, trial condition (i.e., visually congruent and static visual), SNR, and choice. Next, using the first N^pca^ principal components (p) of matrix **M** that explained >80% of the variance, we generated a “denoised” population response: **M**_pca_=**DM**, where 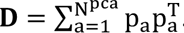.

As a function of each neuron, we computed a time-dependent, multivariate linear regression for each spiking neuron that identified how each task variable modulated neural activity. The estimated regression coefficients were: 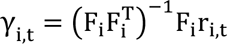. The first n rows of F contained the trial-by-trial values of the n task variables (e.g., trial condition); the number of columns of F_0_ was equal to the number of trials. γ can be considered the direction in a state space that accounts for the variance associated with a particular task variable.

Like with **M**, we denoised the regression coefficients: γ^23+^ = **D**γ. Next, we found the time t_max_ that corresponded to argmax_(_|γ^23+^|, which yielded γ^4+5^. Finally, we found the orthogonal axis of each task variable by orthogonalizing **G^max^** by a QR decomposition: **G^max^**=**QR**. **G^max^** is the set of γ^4+5^ corresponding to each task variable, **Q** is an orthogonal matrix, and **R** is an upper triangular matrix. Finally, p_6,3_ is the task-modulated neural trajectory: 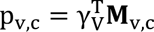. **M** was reorganized as a function of each condition c; for example, each c is one of values indexed by the variable SNR.

### Data and code availability

Data are available on request, and the code on available on github (https://github.com/HuaizhenC/multisensory-effect-in-AC.git).

## Acknowledgements

CH, HS, and YEC were supported by grants from the National Institutes of Health and the Army Research Laboratory. MAE was supported by a grant from the National Institutes of Health.

## Author Contributions

HC conceptualized the study, conducted the behavioral and neurophysiological experiments, analyzed the data, and wrote the paper. HS conducted the behavioral training and behavioral experiments. MAE analyzed the data and wrote the paper. YEC conceptualized the study, analyzed the data, and wrote the paper.

## Declaration of interest

The authors declare no competing financial interests.

## Notes

### Competing Interest Statement

The authors have declared no competing interest.

